# Dopamine neurons change the type of excitability in response to stimuli

**DOI:** 10.1101/053637

**Authors:** Ekaterina Morozova, Denis Zakharov, Boris Gutkin, Christopher Lapish, Alexey Kuznetsov

## Abstract

The dynamics of neural excitability determine the neuronal response to stimuli, its synchronization and resonance properties and, ultimately, the computations it performs in the brain. We investigated the dynamical mechanisms underlying the excitability type of dopamine (DA) neurons, using a conductance based biophysical model, and its regulation by intrinsic and synaptic currents. By calibrating the model to reproduce low frequency tonic firing, NMDA excitation is balanced by GABA-mediated inhibition and leads to type I excitable behavior characterized by a continuous decrease in firing frequency in response to hyperpolarizing currents. Furthermore, we analyzed how excitability type of the DA neuron model is influenced by changes in the intrinsic current composition. A subthreshold sodium current is necessary for a continuous frequency decrease during application of a negative current, and the low-frequency balanced state during simultaneous activation of NMDA and GABA receptors. Blocking this current switches the neuron to type II. Enhancing the anomalous rectifier Ih current also switches the excitability to type II. Key characteristics of synaptic conductances that may be observed in vivo also change the type of excitability: a depolarized GABAR reversal potential or co-activation of AMPARs leads to an abrupt frequency drop to zero, which is typical for type II excitability. Coactivation of NMDARs together with AMPARs and GABARs shifts the the type I/II boundary toward more hyperpolarized GABAR reversal potentials. To better understand how altering each of the aforementioned currents leads to changes in excitability profile of DA neuron, we provide a thorough dynamical analysis. Collectively, these results imply that type I excitability in dopamine neurons might be important for low firing rates and fine-tuning basal dopamine levels, while switching excitability to type II during NMDAR and AMPAR activation may facilitate a transient increase in dopamine concentration, as type II neurons are more amenable to synchronization.

**Author summary:** Dopamine neurons play a central role in guiding motivated behaviors. However, complete understanding of computations these neurons perform to encode rewarding and salient stimuli is still forthcoming. Network connectivity influences neural responses to stimuli but so do intrinsic excitability properties of individual neurons, as they define their synchronization properties and neural coding strategy. We investigated the excitability type of the DA neuron and found that, depending on the synaptic and intrinsic current composition, DA neurons can switch from type I to type II excitability. In short, without synaptic inputs or under balanced excitatory and inhibitory inputs DA neurons exhibits type I excitability, while excitatory AMPAR inputs can switch the neuron to type II. Type I neurons are best suited for coding the stimulus intensity due to their ability to smoothly decrease the firing rate. Type I excitability might be important for achieving low a basal DA concentration necessary for normal brain functioning. Switching to type II excitability further enables robust transient DA release of heterogeneous DA neuron population in response to correlated inputs, partially due to evoked population synchrony.

## Introduction

Midbrain dopamine (DA) neurons predominantly fire in a low frequency, metronomic manner (i.e. tonic) and display occasional, yet functionally important, high frequency, burst-like episodes [1,2]. While tonic firing is observed in isolated preparations (i.e. slices), it is also observed in vivo, where active synaptic inputs make the firing pattern more variable [3,4]. Tonic activity is important for maintaining a constant basal level of dopamine in projection areas. Accordingly, abnormal basal DA levels are linked to psychiatric disorders from depression to schizophrenia [5,6]. While the maintenance of basal DA levels seem to be critical for normal brain function, a consistent picture has not yet emerged regarding how changes in firing patterns of the DA neuron facilitates this important biological function.

Background activity of the DA neuron appears to rely on the intrinsic pacemaking mechanism that generates tonic firing. Based on experimental identification of ion channels [7–18] and modeling studies [4,19,20] the maintenance of tonic firing has been shown to rely on the interactions of the voltage gated Ca^2+^ and SK-type Ca^2+^-dependent K^+^ currents. This interaction periodically brings the neuron to the spike threshold and generates a metronomic firing pattern. In contrast, spike-producing currents (fast sodium and the delayed rectifier potassium) play a mostly subordinate role in this dynamic, adding a spike on top of the oscillations without significant changes to the period or shape of voltage and calcium oscillations [4]. This mechanism is called a subthreshold Ca^2+^-K^+^ oscillatory mechanism. The specific composition of currents contributing to oscillations determines the response of the DA neurons to stimuli, their synchronization properties and, ultimately, the computations they perform. In this paper, we use recent experiments to calibrate the dynamical properties of the DA neuron and determine its excitability type.

Type II neurons, such as inhibitory interneurons neurons in the cortex, display precise spike timing even in the presence of noise, and therefore suitable for the implementation of spike time coding [21,22]. A type I neuron, such as a weakly adapting cortical pyramidal neuron, was shown to relay the stimulus rate by modulating its own frequency, and, therefore, represented rate coding [22]. Further, type II neurons display resonance and controlled synchronization in a network of neurons [22,23,24]. A standard method to classify neuronal excitability is via characterizing the frequency-to-input relationship, or F-I curve. Two major types of excitability can be determined based on how the onset of tonic firing occurs as the applied current increases and the neuron is released from quiescence at the hyperpolarized rest state [25]. A type I-excitable neuron can fire at an arbitrary low frequency near the onset of firing, whereas a type II neuron shows a discontinuous jump to a minimal frequency above a certain current threshold and fires only in a limited range of frequencies [21]. For neurons that are tonically active without any injected current, for example DA neurons, the transition to the non-spiking rest state occurs when a sufficiently strong hyperpolarizing current is injected. For these neurons, the excitability type would be defined by the transition from tonically firing to quiescent/excitable: again type I would show a smooth frequency decrease to zero, while type II should show an abrupt transition to quiescence. At this point, there is no direct evidence defining to what type of excitability the DA neurons belong, and how the different intrinsic conductances and the different synaptic inputs influence its type. Information about the type of excitability will allow us to predict the behavior of the DA neuron during application/blockade of different currents and better understand computations it performs in different input conditions (e.g. rate coding vs. resonance at a particular input frequency).

A number of experimental studies provide indirect evidence of excitability type of the neuron in control and during activation of synaptic inputs. It has been shown that the firing rate of DA neuron increases linearly in response to a ramping depolarizing current until it goes into depolarization block (e.g. [3,26]). Further, injection of a tonic hyperpolarizing current to the regularly firing DA neuron in vitro increases its interspike intervals [7]. The firing properties of the neuron in response to a combination of tonic inhibitory and excitatory synaptic conductances were investigated by Lobb and colleagues [27,28]. By a dynamic clamp technique, they injected an inhibitory γ-Aminobutyric acid (GABA) and an excitatory *N*-methyl-D-aspartate (NMDA) receptor conductances in SNc DA neurons. Injection of tonic GABAR conductance decreases the firing rate of the neuron several-fold. Furthermore, the neuron fired at low frequencies when NMDAR and GABAR conductances balanced each other. Thus, NMDAR activation, which strongly increases the firing frequency [29–32], can be effectively compensated by GABAR activation. Such compensation would be impossible if the inhibition produced an abrupt transition to quiescence and the neuron jumped from a high frequency to zero. The compensation suggests, again, a smooth frequency decrease upon GABAR activation rather than an abrupt transition to the rest state at hyperpolarized potentials. Together, these data resemble the tonic firing/quiescence transition in type I neurons with two distinctions. First, the transition parameter is not an injected current, but an ohmic GABAR conductance. In experiments, a conductance has already been used instead of an injected current to determine the neuronal excitability [33]. Second, the co-activation of the NMDA receptor introduces an additional parameter (its maximal conductance). Both of these extend the definition of excitability into the space of synaptic conductances. Formally, the excitability type is an intrinsic property of a neuron, yet viewing synaptic inputs as changing excitability of a neuron is a powerful concept used to understand neuron dynamics *in vivo* [22,34]. We used the experiments described above to parameterize a model of the DA neuron, determine its type of excitability, and determine how intrinsic and synaptic currents shape the excitability type and, therefore, the computational properties of the neuron.

These experiments suggest that the DA neuron exhibits type I excitability in isolation from synaptic inputs and under the balanced influence of excitatory and inhibitory synaptic conductances. However, the excitability type has been shown to vary depending on the intrinsic currents and network connectivity [34,35]. For example, modeling results suggest that changes in the intrinsic currents, e.g. L-type Ca^2+^ current, can switch excitability type of the DA neuron [36,37]. Further, it has been observed that SNc DA neurons exhibit subthreshold resonance [38], which is associated with type II excitability, and the SK and the Ih current play a role in generating the resonance. Here we address the seemingly incompatible variability in the experimental results under different conditions by studying the contribution of intrinsic and synaptic currents to regulation of the low-frequency DA neuron firing.

## Results

### Balance of tonic NMDA and GABA inputs

Our first goal was to reproduce the experimentally-observed compensatory influence of NMDAR and GABAR conductances [27]. Using the dynamic clamp technique, it was shown in vitro that a balanced injection of GABAR and NMDAR conductances leads to DA neurons firing at frequencies comparable with background frequencies (1-5 Hz). Removal of inhibition in such conditions evokes a disinhibition burst. Figure 1A reproduces the voltage traces obtained in the experiments by Lobb et al. 2010 [27]. The simulated DA neuron is tonically active at 1.5 Hz during tonic co-activation of NMDA and GABA receptors (*g*_*NMDA*_ =16.9mS/ cm^2^, *g*_*GABA*_ = 5*mS*/ cm^2^). Removal of the GABAR conductance produces an episode of high-frequency firing (Fig. 1 C). Removal of the NMDAR conductance produces a pause in firing (Fig. 1 C).

**Figure 1:**
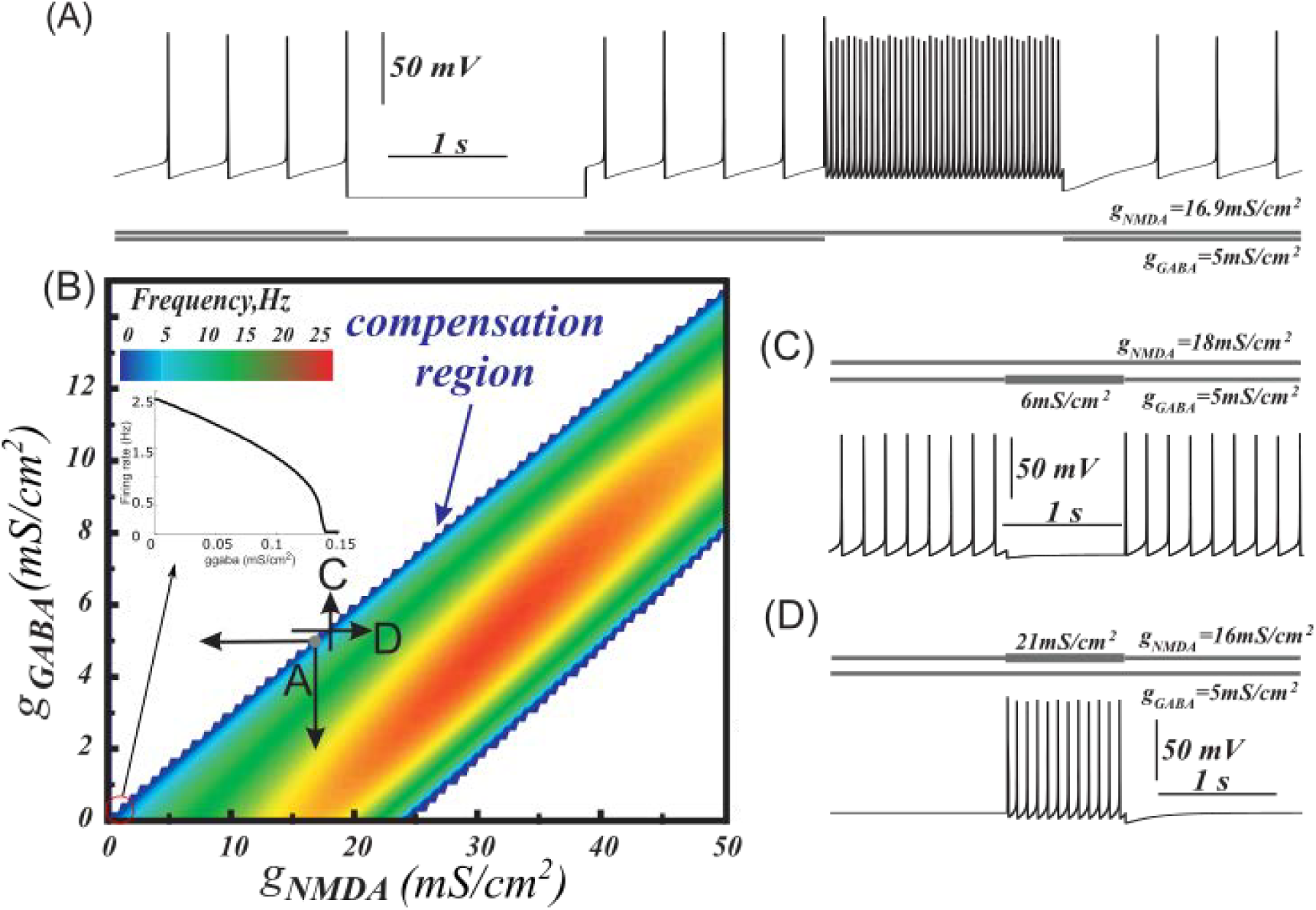
Balance of NMDA and GABA receptor activation and two ways of eliciting bursts or pauses of the DA neuron. (A) Tonic co-activation of NMDA and GABA receptors balance each other and supports low frequency firing in the DA neuron model. Transient deactivation of NMDAR produces a pause, whereas deactivation of GABAR results in a high-frequency burst. (B) Heat plot of the frequency distribution on the plane of NMDAR and GABAR conductances. (C) As an alternative to the deactivation of NMDAR in (A), a pause may be produced by further activation of GABAR. (D) Strengthening NMDA excitation produces a burst, even if GABAR activation initially blocks firing.

We explored the range of NMDAR and GABAR conductances that produce tonic firing in the DA neuron model (Fig. 1 B). Compensation of NMDAR and GABAR activation can be readily achieved near the upper boundary of the firing region (Fig. 1 B, blue). When both receptors are activated, low frequency tonic activity is observed (Fig. 1 A). The dot labeled as A on the heat plot indicates conductances taken for this simulation. As in the experiments [27], the balanced region is stretched linearly on the conductance plane with NMDA/GABA slope around 3.4. Moving to the left on the diagram corresponds to deactivation of the NMDAR current and blocks DA neuron firing due to the remaining GABAR activation (Fig. 1 A). A pause may also be produced by stronger activation of the GABAR (Fig. 1 C). Conversely, moving down on the diagram corresponds to deactivation of the GABAR and evokes high-frequency firing (Fig. 1 A). The firing frequency also increases by moving from the upper boundary of the firing region to the right (increasing NMDAR conductance; Fig. 1 C). These two directions correspond to two ways of eliciting a DA neuron burst: strengthening NMDA excitation or removing inhibition respectively.

Excessive tonic NMDAR activation leads to a depolarization block, as shown in Fig. 1B at high NMDA and low GABA receptor conductances. Interestingly, application of a tonic GABA conductance in combination with an excessive NMDA conductance may rescue high-frequency firing in the model (Fig. 1 B). Thus, the compensatory influence of GABAR activation removes depolarization block induced by an excessive NMDAR activation and restores the intrinsic oscillatory mechanism required for tonic firing.

### Compensatory action of asynchronous NMDA and GABA inputs

We next considered how our model would behave when the inputs display a temporal structure reflective of in vivo conditions. Particularly, we studied the effect of irregular asynchronous GABA and Glu synaptic inputs on DA neuron activity. We quantified changes in firing rate and regularity of DA neuron in response to synaptic inputs of different strengths (Fig. 2 A 1-2, see methods for the detailed description of how these inputs were produced). Similar to the case of tonic inputs, we identified a parameter region where the activity of the excitatory and inhibitory inputs balance to produce low frequency DA neuron firing at rates similar to background firing (Fig. 2 A 1, between the black lines). This happens because asynchronous GABA and Glu inputs (see rasters in Fig. 2 B 1, 2) activate GABARs and NMDARs nearly tonically (Fig. 2 B 3, 4) and provide quasi-constant levels of inhibition and excitation to the DA neuron respectively. Under the influence of these two inputs, the DA neuron fires similarly to the *in vitro-*like conditions (tonic inputs), but with less regularity, which is typical of the background firing *in vivo*. An example voltage trace of the DA neuron in response to synaptic inputs formed by asynchronous Glu and GABA populations is shown in Fig. 2 B 5. Fluctuations in the firing of neural populations innervating the DA neuron can produce irregular firing as observed in *in vivo* experiments.

**Figure 2:**
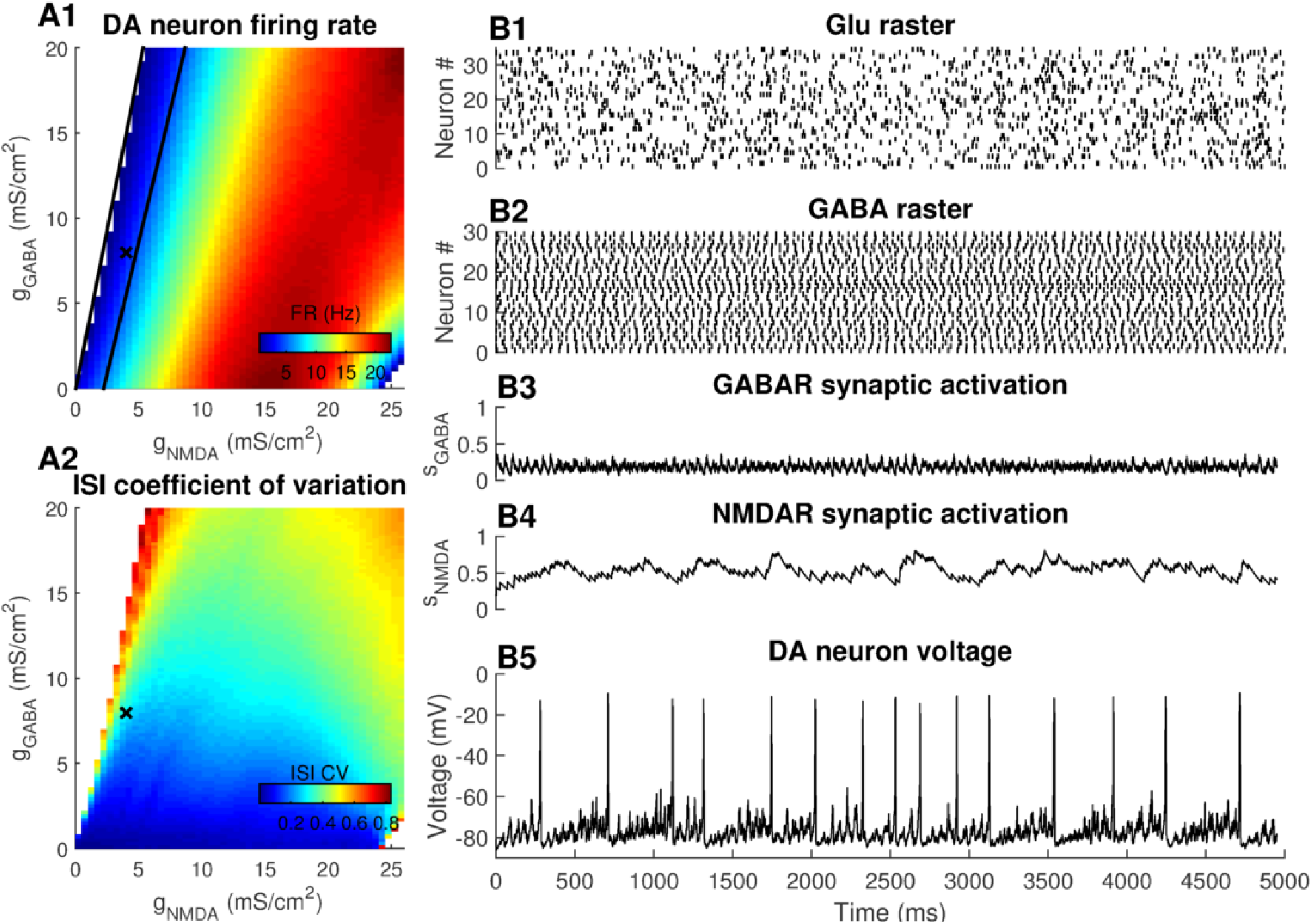
Firing rate and regularity of the DA neuron firing receiving asynchronous synaptic Glu and GABA inputs. (A1) Firing rate. Balanced activity of Glu and GABA populations results in low-frequency DA neuron firing (between black lines) (A2) Coefficient of variation (CV) of the ISI. For majority of *g*_*NMDA*_, *g*_*GABA*_ values, CV<0.5, indicating low variability in DA neuron firing. (B 1-5) Balanced asynchronous GABA and Glu inputs provide constant noisy levels of inhibition and excitation respectively and result in DA firing rates similar to the background firing rates.

### DA neuron excitability under control and during balanced input conditions: type I

The smooth frequency decrease to zero as the neuron transition to quiescence when GABAR conductance increases suggests type I excitability for the DA neuron both in *in vitro* and *in vivo* like cases (Fig. 1 B, 2 A 1). However, there is a difference between the standard definition of excitability type and the case we consider here: excitability is classically defined by the structure of the transition between spiking and hyperpolarized rest state induced by an injected current, as opposed to a synaptic conductance. To highlight this difference, we plot the frequency of the DA neuron as a function of the negative applied current (Fig. 3 A inset) instead of the GABAR conductance (as in Fig. 1A 1 inset). We can see that indeed our DA neuron model is type I under the standard definition with a continuous f-I curve. Further, we extend the dependence into a 2-dimensional heat plot (Fig. 3A main), where vertical axis is the hyperpolarizing current and the horizontal axis is NMDAR conductance as in Fig. 1B. As expected from the dependence on the GABAR conductance (Fig. 1A 1 inset), the frequency smoothly decreases to zero as a stronger hyperpolarizing current (negative) is injected. The similarity reflects that the increase of GABAR conductance increases the voltage-independent current given by *g*_*GABA*_*E*_*GABA*_ in (9), which is negative and equivalent to a hyperpolarizing injected current. Interestingly, the frequency dependence on hyperpolarizing current becomes steeper and the transition becomes more abrupt at higher NMDAR conductance, and the slope of the boundary increases (Fig. 3 A). Due to the greater slope of the boundary, the firing region narrows as the hyperpolarizing injected current grows, and the high-frequency firing can no longer be achieved. This is different from the results shown in Fig. 1B because, as the GABAR conductance increases, the ohmic part, *g*_*GABA*_*v*, induces a significant difference between the synaptic and injected currents. It is interesting to note that the slope of the boundary increases not due to a change in the type of excitability, but because the rest state under a hyperpolarizing current injection is formed at lower voltages (Fig. 3 B 3), where the NMDAR conductance (10) shuts off due to its magnesium block. By contrast, in the case of GABAR input (Fig. 1), the rest state cannot emerge at voltages that are below the GABAR reversal potential. Thus, we predict that DA neurons display the properties of type I excitability in control and in the balanced state. We further investigate the influence of intrinsic and synaptic currents on the excitability type of the DA neuron and relate these results with the type of transition to the hyperpolarized quiescence.

**Figure 3:**
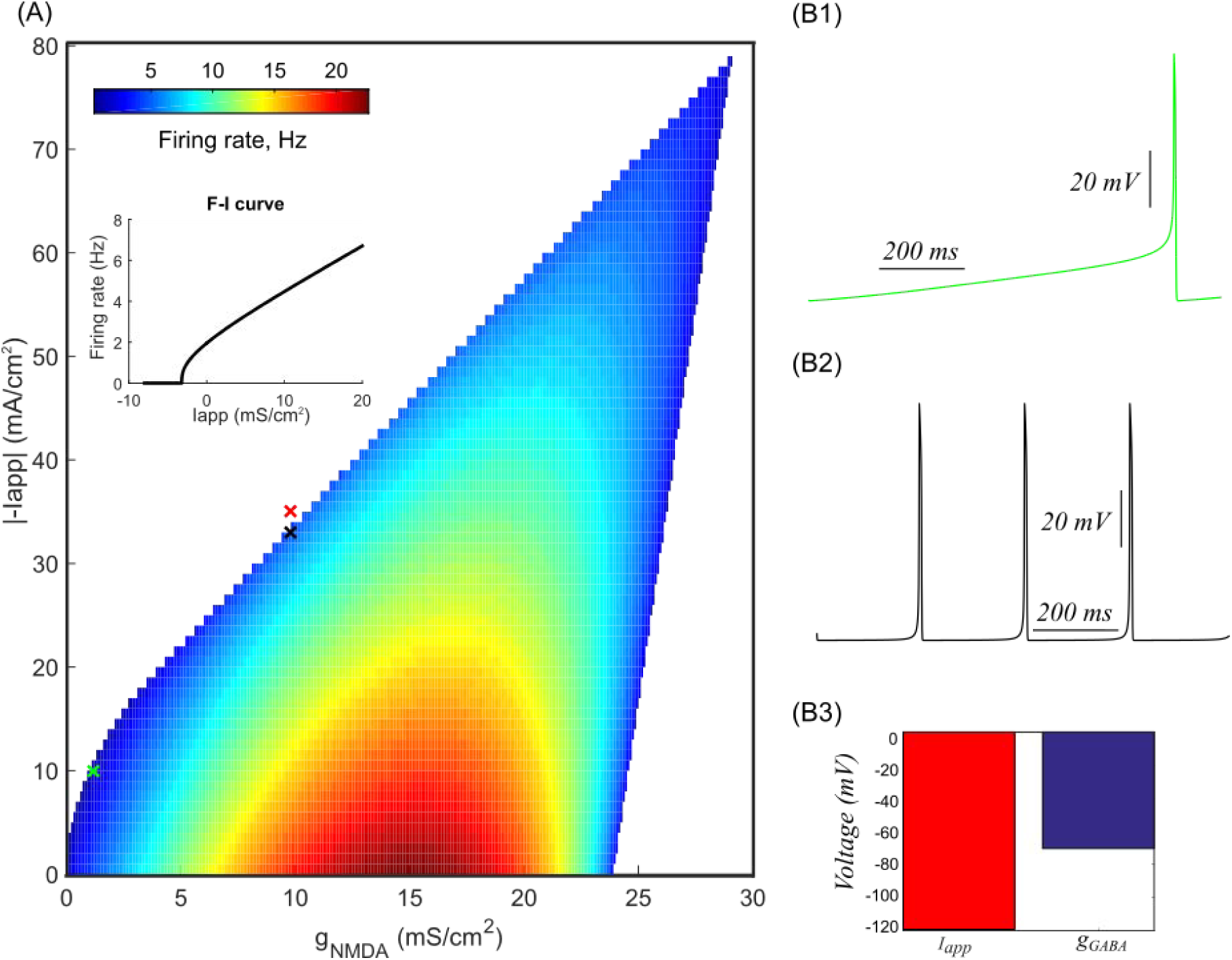
Smooth transition to zero frequency during application of hyperpolarizing current suggestis type I excitability of the DA neuron. (A) The heat plot of the frequency distribution on the plane of NMDAR conductance and hyperpolarizing applied current for the model of the DA neuron. Note that the vertical axis shows the absolute value of the current. The transition to zero frequency remains smooth, which indicates type I excitability. However, the transition becomes steeper at greater hyperpolarizing applied currents. The insert illustrates a smooth frequency decrease during application of the negative current. (B 1-2) Two examples of the voltage traces corresponding to the parameters shown by red and black crosses in panel A. (B3) The rest state is formed at lower voltages in case of negative applied current (red bar) than in case of applied GABAR conductance (magenta bar). The value of NMDAR conductance is the same for both cases and indicated by a red cross in panel (A).

### Influence of intrinsic currents on the type of excitability

*The role of Ca*^*2*+^ *and Ca*^*2*+^-*dependent K*^+^ *currents*

The subthreshold Ca^2+^-K^+^ oscillatory mechanism underlies the generation of low frequency background firing in the majority of DA neurons. We found that, if considered in isolation, the Ca^2+^-K^+^ oscillatory mechanism provides type II excitability, which is incompatible with the experiments reproduced above. In order to study this we use a reduced scenario in our model, where we block the subthreshold sodium current to isolate the Ca^2+^-K^+^ oscillatory mechanism. The interaction of the L-type voltage-gated Ca^2+^ and the calcium-dependent potassium currents periodically brings the neuron to the spike threshold and generates a metronomic firing activity pattern. Application of an inhibitory input (GABA or hyperpolarizing current) to this model gradually reduces the amplitude of voltage oscillations instead of decreasing the firing frequency gradually (see nullcline analysis below in Fig. 6 A). This transition is typical for systems where the oscillations are terminated via an Andronov-Hopf bifurcation: the oscillatory trajectory (limit cycle) decreases in amplitude and merges with an equilibrium state. An abrupt transition to zero frequency quiescence upon hyperpolarization suggests a type II excitability of the neuron. Thus, these currents are important for producing metronomic firing, but other intrinsic currents allow for a gradual frequency decrease during application of a hyperpolarizing input observed experimentally.

**Figure 6:**
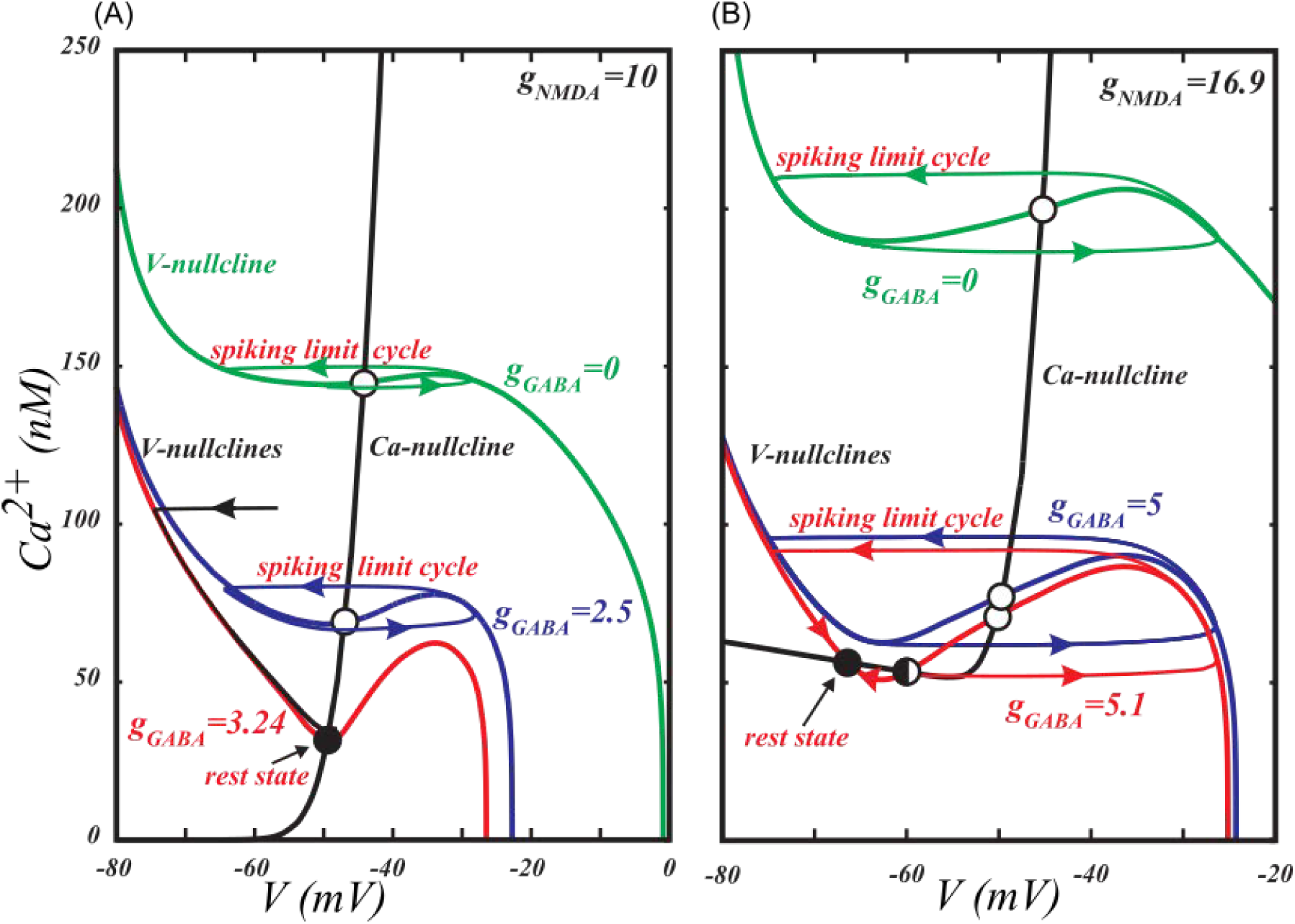
The subthreshold sodium current changes the type of transition to hyperpolarized rest state induced by GABAR activation. In both cases the growing GABAR conductance leads to a downward shift of the voltage nullcline (solid folded curve). (A) In the pure Ca^2+^-K^+^ mechanism for voltage oscillations, inhibition leads to a transition to the rest state through an Andronov-Hopf bifurcation, which occurs with little change in the firing frequency. The Andronov-Hopf bifurcation is defined as the disappearance of a closed trajectory representing firing (limit cycle) by shrinking in amplitude and merging with an equilibrium state. (B) If the Ca^2+^-K^+^ mechanism is augmented by a subthreshold Na^+^ current, the transition occurs through a Saddle-Node on Invariant Circle bifurcation (SNIC), which corresponds to a gradual decrease in the frequency to zero. The SNIC bifurcation is defined as the emergence of new equilibrium states that interrupt the limit cycle.

*The role of a subthreshold sodium current*.

We found that the subthreshold sodium current is necessary for gradual frequency deceases during application of a hyperpolarizing input, as well for a frequency range that spans the observed control frequencies of the DA neurons during a balanced tonic NMDAR and GABAR activation. The reduced firing frequency in the balanced state comes about because the inputs create a slow “bottleneck” effect in the subthreshold voltage range where the hyperpolarizing inputs nearly cancel the depolarizing ones (Fig. 4, also see methods for a more detailed description) and expand the interspike interval. This input balance is achieved due to the contribution of the subthreshold sodium current into the pacemaking mechanism of the DA neuron. By contrast, in the reduced model that includes only Ca^2+^ and Ca^2+^-dependent K^+^ currents into the mechanism, the inhibitory input cannot restore appropriate frequency, but instead blocks the voltage oscillations. The inclusion of the subthreshold sodium current allows the firing frequency to vary without compromising the oscillatory mechanism. It allows us to reduce the frequency to an arbitrary low value upon application of a hyperpolarizing input and thus leads to type I excitability of the DA neuron.

**Figure 4:**
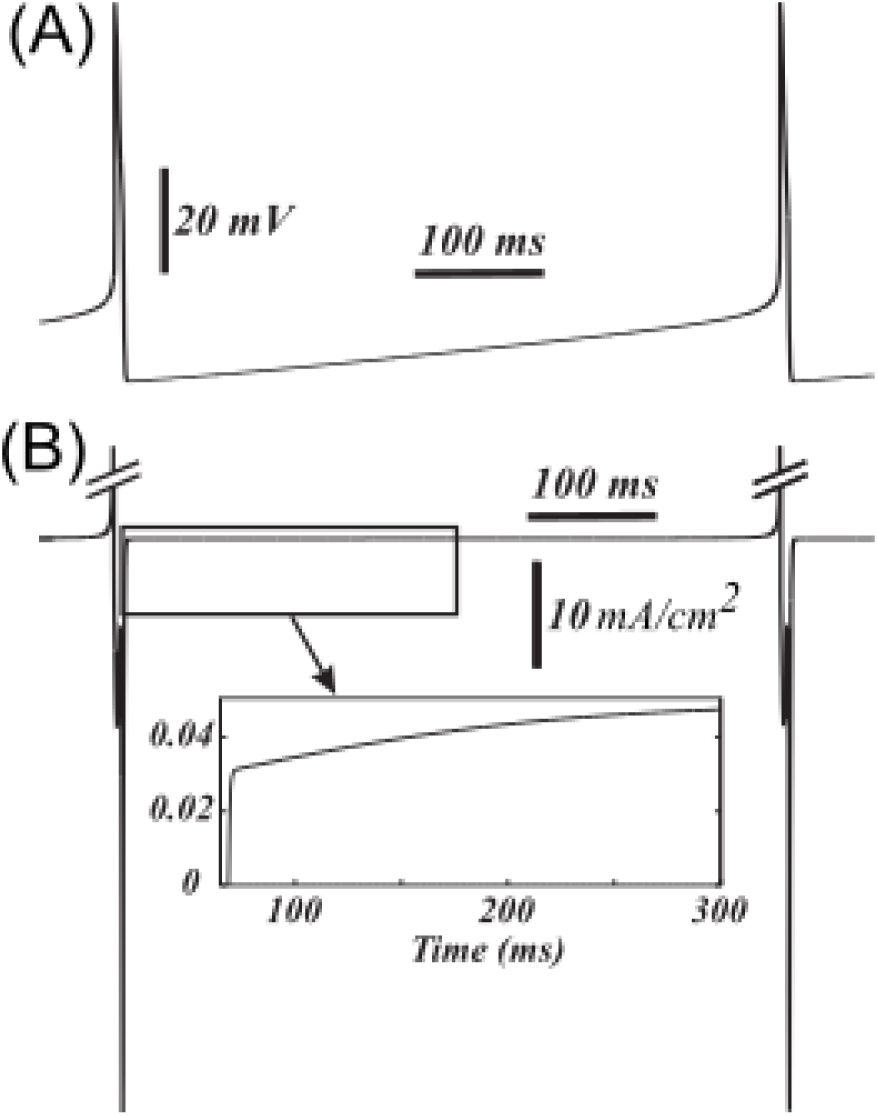
The time series of the total transmembrane current (B) shows near cancellation of the current during the interspike interval, which creates a bottleneck effect. The voltage is shown in (A) for a reference.

### A mechanism for NMDA-GABA balance: SNIC bifurcation

Mathematical analysis allows for a better explanation of how the subthreshold sodium current changes the response of the neuron to a combination of excitatory and inhibitory inputs. First, we further reduce the model by removing the fast sodium and the delayed rectifier potassium spike-producing currents. We now define the model as having fired a spike when the voltage crosses a putative spike-threshold (set at −40 mV). Figure 5 shows that the frequencies displayed by the model do not significantly change. This reproduces the experiments, in which blockade of the spike-producing currents do not significantly change either the frequency of background oscillations [4] or the NMDA-evoked high frequencies [32]. The reduction decreases the number of variables in the model and enables standard nullcline analysis because it applies only to two-dimensional systems [39].

**Figure 5:**
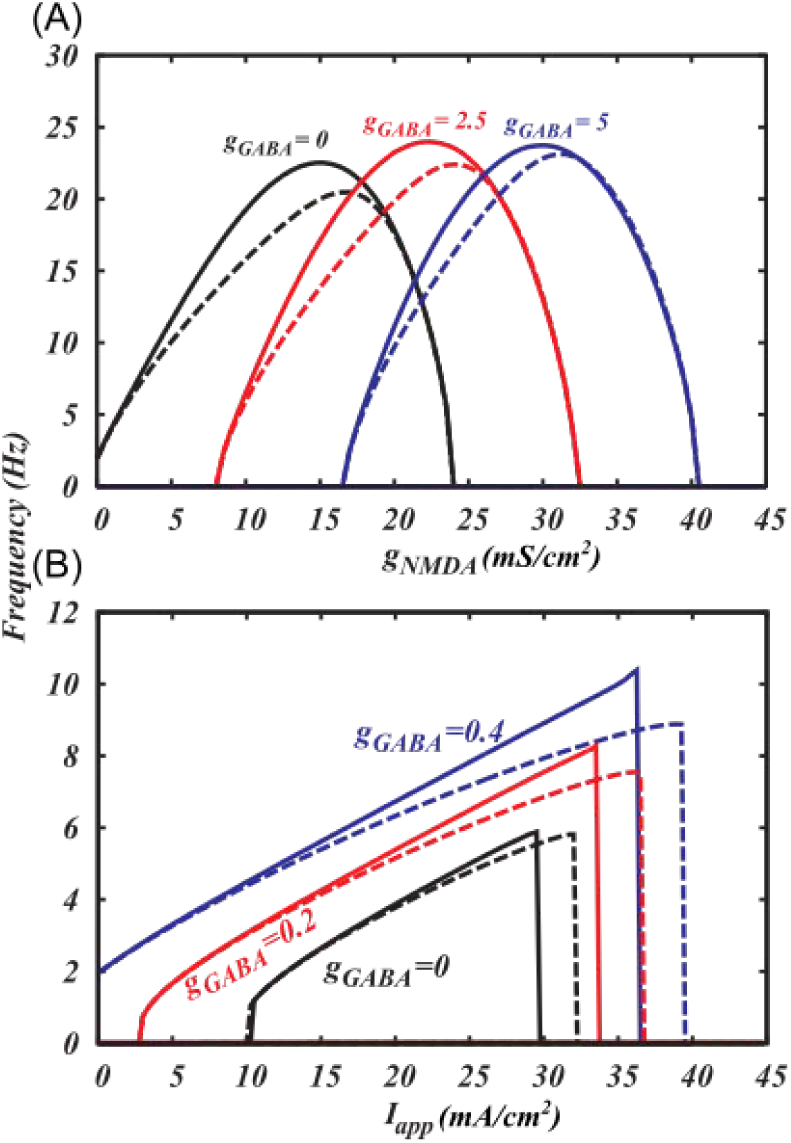
Spike-producing currents (I_Na_ and I_DR_) have no significant influence on the frequency growth induced by NMDA receptor activation (A) or an injected current (B). The frequencies are measured in the model with (dashed) and without (solid) these currents.

Our reduced Ca^2+^-K^+^ model for voltage oscillations (e.g. [4]) shows that the transition to quiescence at a hyperpolarized voltage occurs via an Andronov-Hopf bifurcation, in which oscillations disappear without a decrease in the frequency. Figure 6 presents standard nullcline analysis of the model, which explains oscillatory behavior and bifurcations mechanistically. In Fig. 6A, the Andronov-Hopf bifurcation occurs as the voltage nullcline shifts down and simultaneously to the right, so that its intersection with the Ca^2+^ nullcline moves across the minimum. In the model with the subthreshold sodium current and Ca^2+^ leak current (Fig. 6 B), the minimum of the voltage nullcline is further away from the steep part of the Ca^2+^ nullcline, so that, when the voltage nullcline shifts down, its minimum touches the flat part of the Ca^2+^nullcline. The proximity of the minimum of the voltage nullcline and the bottom part of the Ca^2+^nullcline creates a “bottleneck” effect: The closer the nullclines, the smaller the vector field (the rate of change) in this neighborhood. The limit cycle is channeled through the gap between the nullclines and, accordingly, the oscillation evolves slowly. In the limiting case, when the minimum of the voltage nullcline touches the bottom part of the Ca^2+^ nullcline, a saddle-node on invariant circle (SNIC) bifurcation occurs: two equilibrium states, a stable (node) and an unstable (saddle) emerge, interrupt the limit cycle, and the period becomes infinite. The closer the bifurcation parameter *g*_*GABA*_ to the bifurcation value, the more time the voltage spends in the bottleneck, creating a long interspike interval. Thus, by introducing the subthreshold sodium current, we change the bifurcation that leads to the quiescence at hyperpolarized potentials from Andronov-Hopf to SNIC.

*The role of Ih current*

DA neurons are often identified by a prominent hyperpolarization-activated cation current (Ih), which gives a voltage “sag” upon injection of a hyperpolarizing current. It has been shown that Ih-expressing DA neurons exhibit smooth frequency decrease pointing to a type I excitability [27,28]. However, another study that investigated the response of the DA neuron to a swept sine wave current, using the impedance (Z) amplitude profile (ZAP) method [38] showed that DA neurons produce a resonance at 2-7 Hz frequency and the resonance is abolished by blocking the Ih current. This suggests that DA neurons are type II excitable and the Ih current plays an important role in producing the resonance. Our model also suggests that DA neurons display type II excitability in the presence of the Ih current. However, at small values of Ih conductance (gh=1-6 mS/cm^2) the model behaves very similarly to the one without the Ih current, which might be the reason of the apparent discrepancy in the excitability type as suggested by experiments. For small values of Ih conductance, the firing frequency of the simulated DA neuron decreases abruptly with the increase in the hyperpolarizing current. However, the discontinuity occurs at very low frequencies (see Fig. 7), which might appear still as a gradual frequency decrease in the experiments. As the strength of Ih conductance increases (gh>6 mS/cm^2) the discontinuity in the F-I curve becomes clearer. This apparent switch in the excitability type might be important in the altered states of the DA system, when the Ih current is potentiated, for example, by EtOH [40,41]. For high conductances of Ih, low frequency tonic firing cannot be produced, which might affect the basal DA levels.

**Figure 7:**
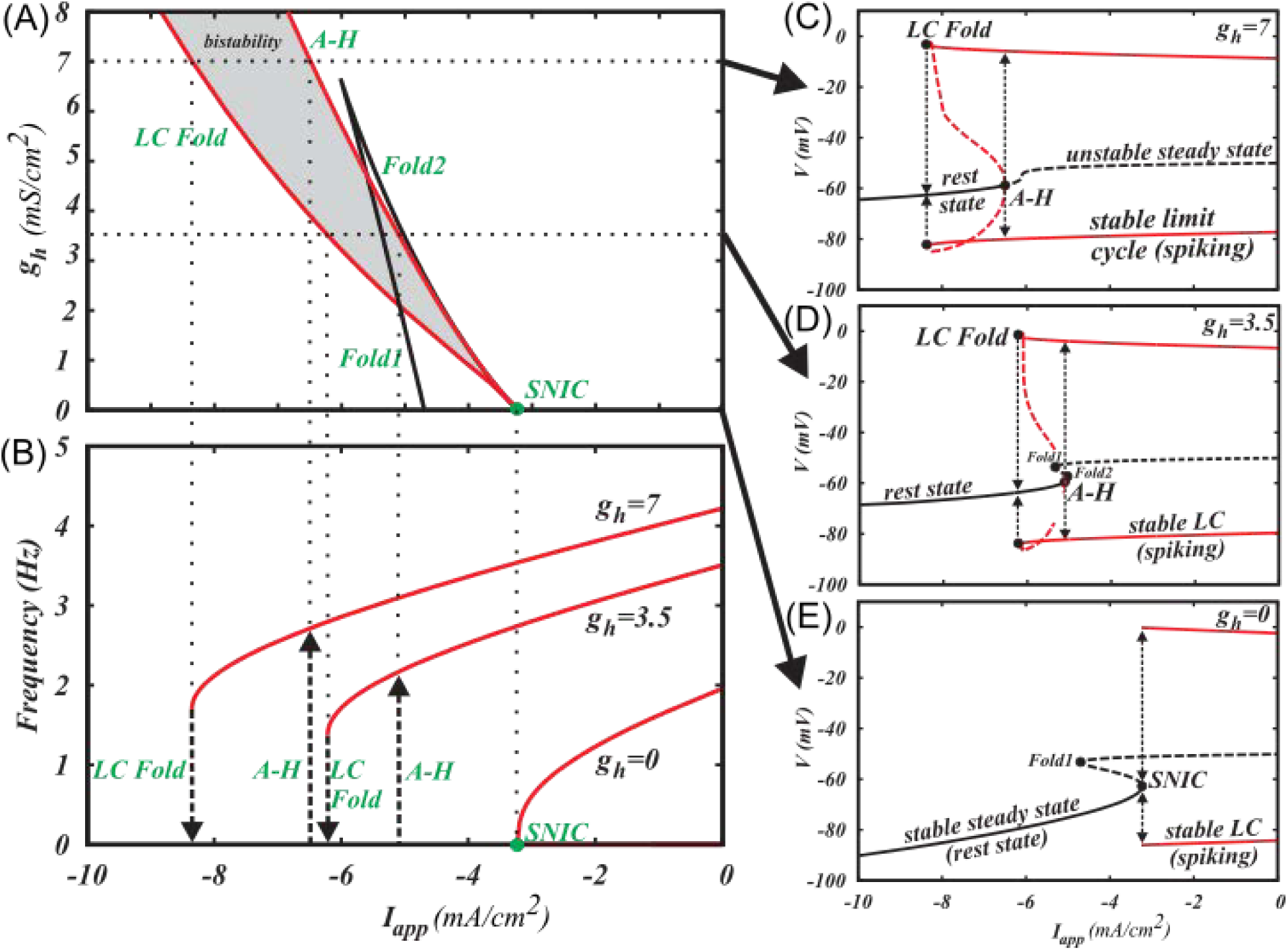
The I_h_ current switches the type of excitability of the DA neuron to type II. (A) Two-parameter bifurcation diagram. See part (C) for explanations. LC Fold is a saddle-node bifurcation of limit cycles, in which two oscillatory solutions, stable and unstable, emerge. A-H is the Andronov-Hopf bifurcation, in which a limit cycle shrinks in amplitude and merges with an equilibrium state. SN is a saddle-node bifurcation of equilibrium states, in which two equilibria (rest states) emerge. SNIC is a saddle-node bifurcation of equilibria on invariant circle, which is the same SN bifurcation that occurs on a limit cycle and, therefore, interrupts it. (B) F-I curves for three values of gh: gh=0mS/cm>2, gh=3.5mS/cm>2 and gh=7mS/cm>2. (C1-3) Representative one-parameter bifurcation diagrams of three different bifurcation scenarios occurring at gh=0mS/cm>2, 0<gh<7mS/cm>2 and gh>=7mS/cm>2. Black curves represent equilibria. Red curves represent minima and maxima along a limit cycle. Solid stands for stable and dashed for unstable solutions. Dotted arrows represent bifurcation transitions. In C1, spiking is blocked as Iapp decreases through the LC Fold bifurcation. If Iapp increases through the same interval, the LC Fold bifurcation stays unnoticed because the equilibrium state remains stable, and only after the A-H bifurcation, spiking emerges. This is called hysteresis, and it creates bistability, in which either spiking or the rest state can be observed depending on the initial conditions for the voltage and Ca^2+^ concentration. In C2, the curve of equilibrium states folds and two saddle-node bifurcations of equilibria occur. The equilibria interrupt the unstable limit cycle (homoclinic bifurcation; not shown), but do not affect the stable limit cycle yet. In C3, the equilibria interrupt the stable limit cycle and destroy it in the SNIC bifurcation.

As we describe earlier, the smooth frequency decrease and the transition to the rest state upon application of hyperpolarizing current or activation of GABAR occurs via a SNIC bifurcation (Fig. 7 C 3). As soon as we include the Ih current into the model, the SNIC bifurcation splits into a saddle-node bifuraction of limit cycles (LC Fold) and two subcritical Andronov-Hopf bifurcations as shown in a two-parameter bifurcation diagram in figure 7 A (see caption for the definitions of bifurcations). Accordingly, the excitability type changes to type II as we introduce Ih. However, for small values of its conductance gh, the two bifurcations are very close together and nearly indistinguishable, thus, the discontinuity in the F-I curve is in a very narrow range close to zero Hz, producing an illusion of a smooth transition to the rest state.

The bifurcation scenario for the values of gh less than 7mS/cm^2 is complex, although this complexity does not affect experimental observations as the affected limit cycle is unstable. As the magnitude of the negative applied current increases, the unstable limit cycle emerging from the Andronov-Hopf bifurcation, disappears as it collides with a saddle equilibrium state in a homoclinic bifurcation (not marked) and then appears again as the saddle disappears (SN1 in fig. 7). The amplitude of the unstable limit cycle grows with the further increase in the negative applied current and, finally, it merges with the stable limit cycle for spiking and annihilates it at the fold bifurcation for limit cycles (LC Fold, Fig. 7 C 2). For higher values of gh>7mS/cm^2 the curve of equilibrium states becomes monotonic, the saddle-node bifurcations of equilibrium states disappear and the transition from spiking to the rest state occurs via the LC fold bifurcation (fig. 7 C 1). Figure 7 illustrates representative examples of one-parameter bifurcation diagrams and F-I curves for all three cases: gh=0, 0<gh<7 and gh>=7.

For a range of negative applied currents, a stable limit cycle coexists with a stable equilibrium state, creating bistability. Thus, depending on the initial conditions, the neuron will be either in a rest (at equilibrium point) or in a repetitive spiking state (at limit cycle). The bistability region grows with the increase in the conductance of the Ih current (Fig 7 A 1). Variations in current strength back and forth across this range will cause the neuron to jump from one state to the other. To check predictions generated by our model regarding the presence of hysteresis with respect to the applied current, slowly rising and falling current ramps can be applied to the DA neuron to switch it from the rest state to the repetitive firing mode and vice versa. The switch points should be different for falling and rising stimuli as shown in figure 7 B. Despite the range of current intensities for coexistence is small relative to the total range for repetitive firing, it affects neuron behavior near the threshold. For example, once the current intensity reaches the value required for the onset of repetitive spiking, small perturbations of current will not switch the neuron back to the rest state. Thus, the presence of hysteresis makes spiking near the threshold more robust.

### Changes in the type of excitability caused by synaptic inputs

*In vivo*, the type of excitability may change due to tonic synaptic inputs [34], and next we explore how this change occurs in the DA neurons. AMPA receptor may co-activate together with NMDA and GABA receptors *in vivo*. By contrast to NMDAR, conductance of which is voltage-dependent, AMPA and GABA receptor currents are purely ohmic. Their combination is also an ohmic current:

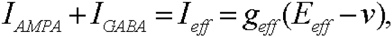

where *g*_*eff*_=*g*_*AMPA*_+*g*_*GABA*_ is a combined synaptic conductance, and

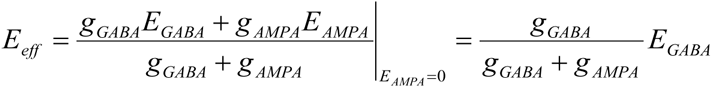

is a synaptic reversal potential. Figure 8 shows the frequency distribution and the type of bifurcation at the transition to the rest state on the plane of these two parameters: conductance and the reversal potential of the ohmic synaptic current. For instance, if the AMPA receptor is blocked, the reversal potential coincides with the GABAR reversal potential, which is in the range of −90 to −70 mV. In this range, an increase in the conductance *E*_*eff*_ leads to a gradual decrease in the frequency to zero and a transition to the rest state via a SNIC bifurcation. By contrast, at higher reversal potentials, the frequency drops to zero abruptly and the transition corresponds to an Andronov-Hopf bifurcation. This suggests a transition to type II excitability for the DA neuron. Thus, elevated GABAR reversal potential or tonic activation of AMPAR leads to a switch in the excitability to type II.

**Figure 8:**
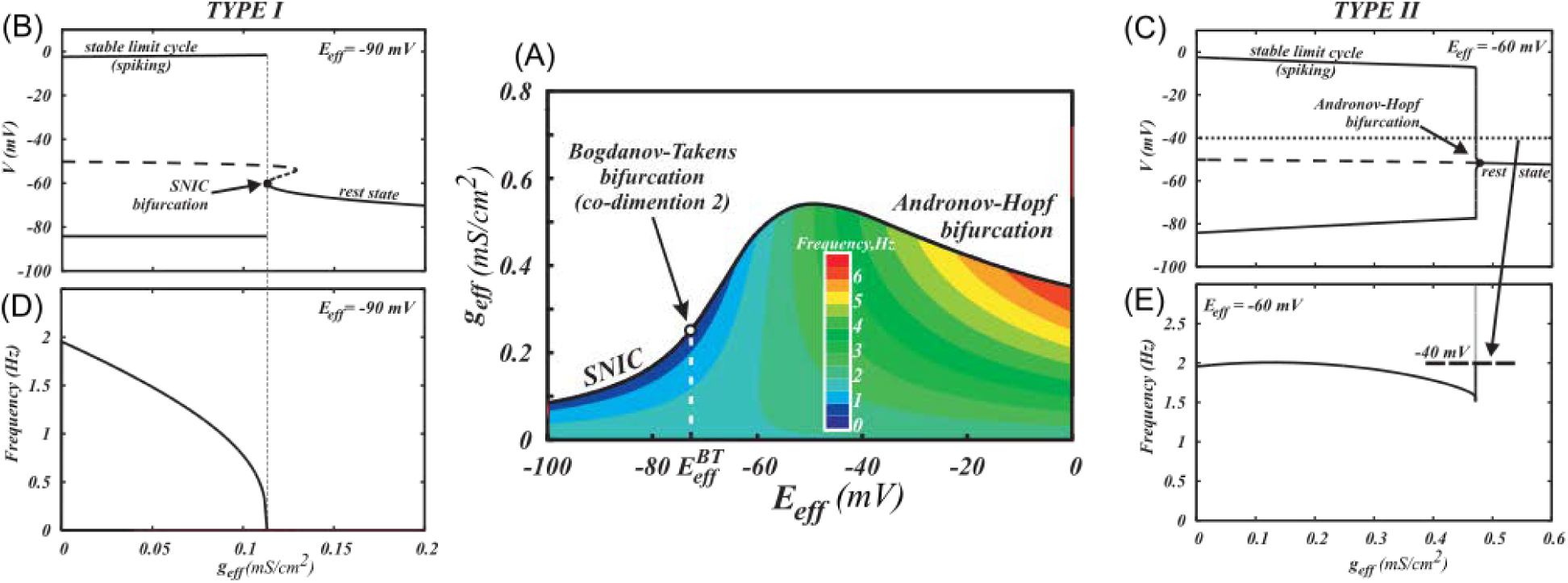
A combinations of ohmic synaptic conductances determines the type of excitability for the neuron. (A) The type of excitability is connected with the type of bifurcation at the transition from spiking to the rest state. The bifurcation type changes at 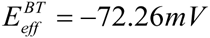, which is called Bogdanov-Tackens point. B) SNIC bifurcation scenario for a low reversal potential of the ohmic synaptic current (*E*_*eff*_= −90*mV*), (D) A smooth frequency decrease to 0 upon application of combined synaptic conductance, corresponding to type I excitability. (C) An andronov-Hopf bifurcation scenario for the higher reversal potential of the ohmic current (*E*_*eff*_ = −60*mV*), (E) An abrupt frequency decrease to 0, corresponding to type II excitability.

### Activation of GABAR cannot rescue firing blocked by tonic AMPA

AMPA quickly induces depolarization block in the DA neurons and firing cannot be restored. The dynamical explanation is that AMPAR activation shifts the minimum of the voltage nullcline across the Ca^2+^ nullcline, so that for high AMPAR conductance values (as well as positive applied currents), voltage oscillations decrease in amplitude and depolarization block occurs (data not shown). Thus, DA neuron firing does not exceed the frequency of ˜10 Hz when driven with AMPAR activation similar to the experimental results (see e.g [29,30,32]), Application of GABA shifts the voltage nullcline even further to the right and makes the stable equilibrium more robust. Therefore, the region of parameters for which spiking is produced is much smaller for combined application of AMPA and GABA, then for combined NMDA and GABA activation (compare Figs. 1 B and 9 A). Therefore, the prediction of our model is that a disinhibition burst can be supported by tonic background activation of NMDA, but not AMPA receptor.

**Figure 9:**
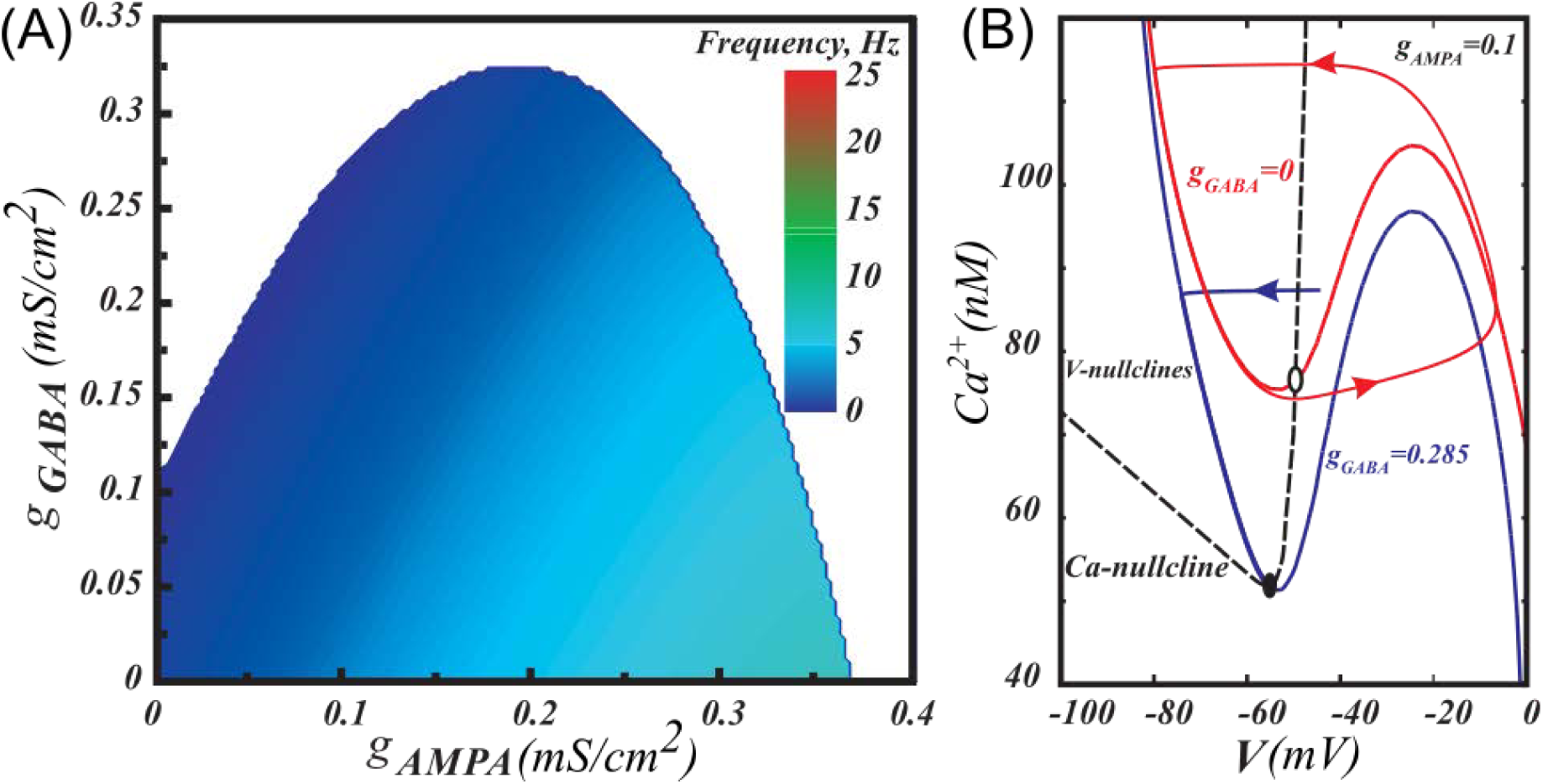
GABA cannot rescue firing blocked by tonic AMPA. (A) Firing persists in a much smaller range of GABAR conductance if co-activated with AMPAR than with NMDAR. (B) The analysis shows that AMPAR conductance transforms the voltage nullcline (solid) differently from NMDAR conductance. This transformation cannot be compensated by GABAR activation.

### Activation of NMDAR shifts the boundary between the excitability types to lower values of the GABA reversal potential

Together with AMPA and GABA receptors, the NMDAR may be also co-activated since glutamate binds to both AMPA and NMDA receptors. To make the analysis of the excitability type possible in the parameter space of all three synaptic currents, we further perform 2-dimensional bifurcation analysis. The point marked Bogdanov-Takens bifurcation in Fig. 8 is a good predictor of the type of excitability at the boundary between spiking and the rest state. Mathematically, it is defined as a junction of the SNIC bifurcation and the Andronov-Hopf bifurcation, as it appears in the figure. In Fig. 10, we plot this point as a function of the NMDA receptor conductance. As in the previous figure, the transition to quiescence occurs as the combined conductance of the ohmic synapses *g*_*eff*_ grows. The information about the specific value of the conductance is omitted in Fig. 10 because the transition occurs in a dimension orthogonal to the plane of the figure. For example, the transition in Fig. 8 is represented by one line at *g*_*NMDA*_=0. The diagram in Fig. 10 shows that the separation between the types of excitability shifts to lower values of the combined reversal potential for the AMPAR and GABAR currents as the NMDAR conductance increases. However, this shift quickly saturates and is restricted to the range of GABAR reversal potentials. Thus, similar to Ih, NMDAR activation may switch the type of excitability of the DA neuron from type I to type II in a certain window of other tonic synaptic currents received by the neuron.

**Figure 10:**
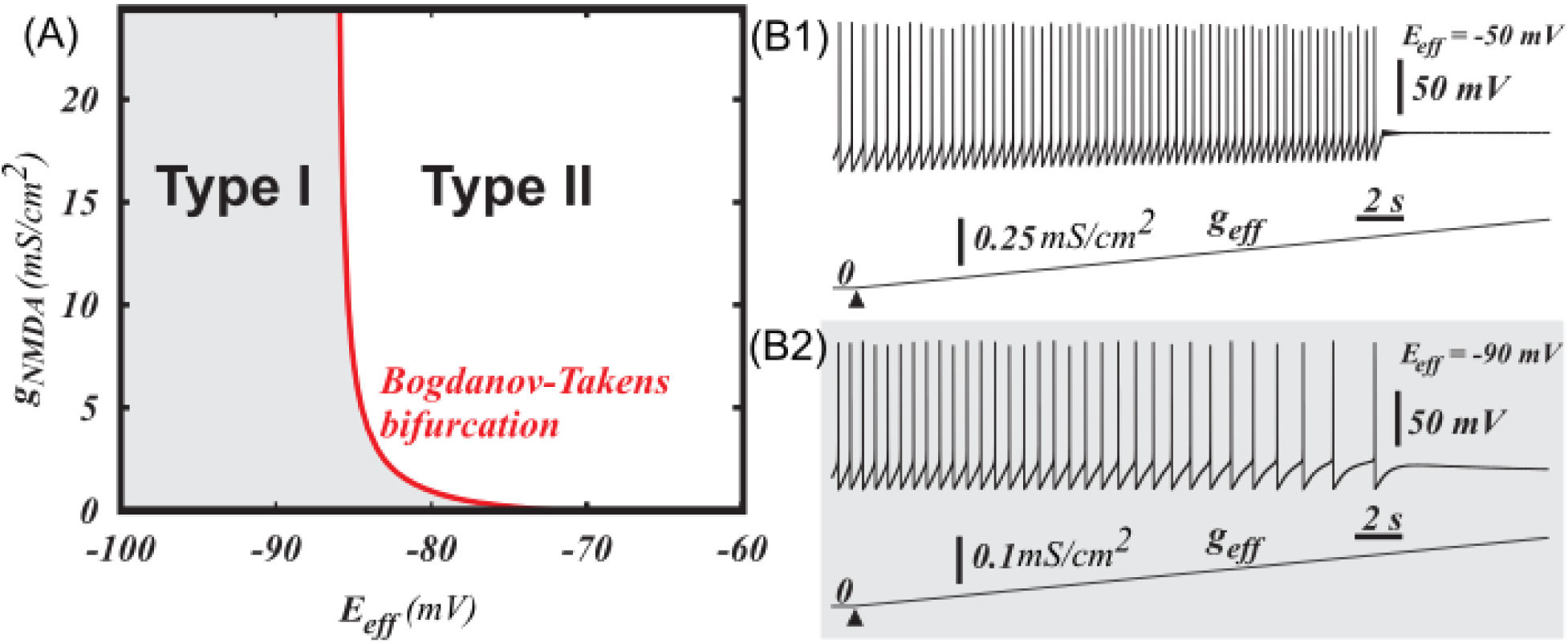
Tonic activation of NMDAR may change the excitability type from the first to the second. (A) The boundary between type I and type II excitability in the DA neuron shifts down with respect to the reversal potential of an ohmic synaptic current as NMDAR conductance grows. (B1-2) Two voltage traces illustrating the excitability types. For any synaptic reversal potential *E*_*eff*_ and NMDAR conductance from the grey region of type I excitability on panel (A), ramping the ohmic synaptic conductance *g*_*eff*_increases the interspike intervals and then blocks firing. For the parameters from type II region, ramping the ohmic synaptic conductance blocks firing without the decrease in the frequency.

### DA release and synchronization in a population of heterogeneous DA neurons is influenced by tonic background synaptic currents

To illustrate the importance of changing DA neuron type of excitability, we simulated heterogeneous populations of DA neurons under two conditions: 1) in control (in the absence of the tonic synaptic inputs), and 2) during the tonic influence of AMPA and GABA inputs. The DA population is electrophysiologically heterogeneous [42–44], and its uncoordinated activity produces a homogeneous low-level DA concentration. In order to have similar firing rates and basal DA levels in both cases, we balanced the increase in firing rate produced by the application of AMPAR conductance with GABAR conductance (note that GABA can balance AMPA only for a very limited range of values). In both cases, DA neurons received correlated fluctuating NMDA inputs (Fig. 11A, see methods for the detailed description of the inputs). Our simulations show that the population of DA neurons that receive the background synaptic tone produces higher DA levels in response to bursty correlated NMDA input than a population without the synaptic tone (Fig. 11 D). As described above, DA neurons are type I excitable in the absence of synaptic tone, while AMPAR activation switches DA neuron excitability to type II. Thus, the transition from type I to type II excitability of the DA neurons is accompanied by higher dopamine release in response to a correlated synaptic input. The higher responsiveness is partially due to a greater synchronization of the DA neurons receiving the synaptic tone, as evident by the higher spikes in their summed activity in Fig. 11C. Type II neurons display more robust synchrony when they receive a common input, even in the presence of independent noise [24,45]. Thus, *in-vivo*background synaptic tone might be important not only for regulating basal DA levels, but also for the responsiveness of the DA neurons, so that they are more ready to produce coincident bursts in response to correlated synaptic inputs.

**Figure 11:**
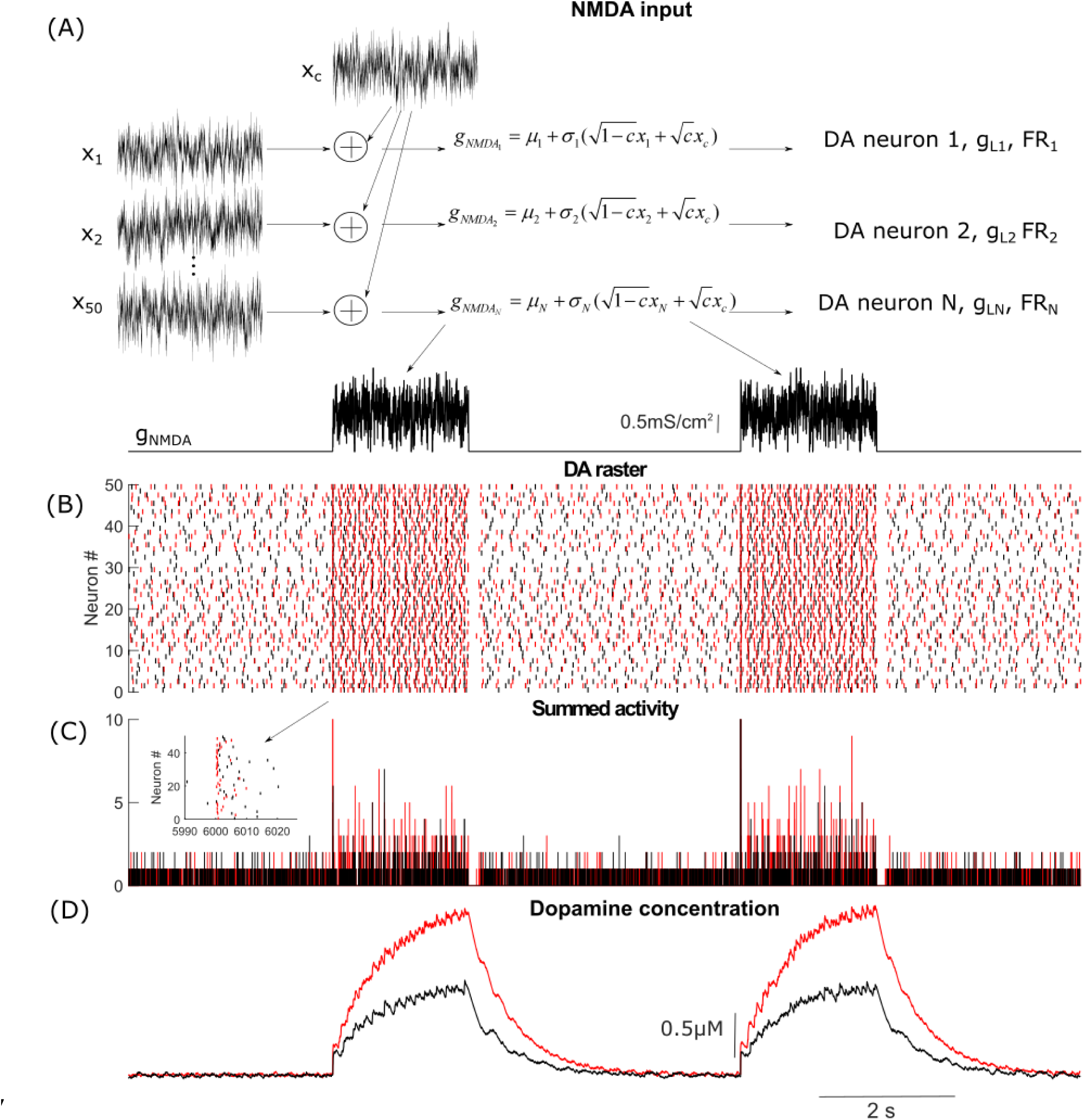
DA release and robustness of synchrony in a population of heterogeneous DA neurons are sensitive to background synaptic tone. (A) Stimulation paradigm in which DA neurons receive correlated fluctuating NMDA synaptic inputs with means of NMDAR conductences *μ*_1_ = *μ*_2_ = *μ*_*N*_ = 1.5*mS* / *cm*^2^ and standart deviations σ_1_ = σ_2_ =…= σ _*N*_ = 0.5*mS* / *cm*^2^. Fluctuations were modeled by Ornstein-Uhlenbeck process with τ = 5*ms* (de la Rocha et al., 2007, Hong et al. 2012). (B) Raster plot showing the firing of two heterogeneous poulations of DA neurons in response to the fluctiating NMDA input. DA neurons shown in black do not receive tonic synaptic inputs, while neurons shown in red receive tonic AMPA and GAB A inputs. The input are balanced such that the firing frequency remains the same. (C) Summed activity of the DA neurons in both populations. The level of synchrony during application of NMDA bursts is higher in the population receiving the tonic synaptic inputs. This is also illustrated in the insert zoomed in on the neural activity at the beginning of the burst. (D) Cummulative synaptic DA concentration produced by activity of the DA population with (red) and without (black) tonic background synaptic inputs. The DA neuron population that recieves tonic inputs releases more dopamine in response to NMDAR stimulation.

## Discussion

### Discussion The excitability type of the DA neuron

The type of excitability for a neuron determines the neurons’ responses to stimuli and their dynamics in a population (phase response curve, synchronization, resonators vs. integrators). The type of bifurcation determines the type of excitability: neural oscillations that arise via an Andronov-Hopf bifurcation have type II excitability, while those appearing via SNIC have type I excitability [39,46]. Based on the bifurcation analysis and frequency responses to hyperpolarizing inputs (negative injected current and GABAR conductance), we have shown that, in control conditions the simulated DA neuron is type I-excitable. It’s known that the type of excitability in vivo may be different from in vitro [34]. In vivo, DA neurons display irregular low-frequency activity occasionally interrupted by high-frequency bursts. This low-frequency regime may reflect the balance of inhibitory and excitatory inputs. We found that, in the most prominent low-frequency regime, the DA neuron is type I excitable, in either high or low synaptic conductance states.

The baseline level of dopamine is very important for the normal function of the brain. The level is determined by the background activity of the DA neurons. This activity is intrinsically generated by the neuron and controlled by its synaptic inputs [47], reviewed in [48]. Thus, the capacity of the DA neuron to adjust its firing rate according to the inputs is vital. The graded response curve of a type I-excitable neuron, as oppose to an on-off response of a type II neuron, provides this capacity. Accordingly, at every level of excitation provided by NMDAR input, inhibitory GABA synaptic conductance can balance it and bring the frequency down to an arbitrary low value. A similar hyperpolarizing current activated by dopamine D2 receptors on the DA neuron has also been shown to slow down its firing rather than abruptly block it all together [49]. These are very important autoregulatory functions of the DA system that allow it to adjust basal DA levels in target areas.

### The role of the subthreshold sodium current

Persistent sodium current is known to amplify subthreshold oscillations (Hu et al, 2002) and increases neural excitability of DA neurons by contributing to spontaneous depolarization in between the spikes [50]. Consistent with experimental observations, the subthreshold sodium current increases the firing rate of the DA neuron in the model. Additionally, we found that the current is necessary for achieving gradual frequency decrease upon application of hyperpolarizing current, thus, maintaining type I excitability of the DA neuron. The type of excitability is determined by the internal properties of the currents contributing to pacemaking in the DA neuron. L-type Ca^2+^ and SK-type Ca^2+^-dependent K^+^ currents are the core currents that traditionally constitute the mechanism [7–18]. However, our model shows that the mechanism results in type II excitability, in which a hyperpolarizing current blocks the voltage oscillations without restoring a low frequency. Our explanation is that, without the inclusion of other currents, the steep part of the Ca^2+^ nullcline is very close to the minimum of the voltage nullcline (Fig. 4 A) because they reflect the same event: opening of the Ca^2+^ channel. This positions the system close to the Andronov-Hopf bifurcation, which occurs whenever the minimum of the voltage nullcline moves across the Ca^2+^ nullcline. When the subthreshold sodium current is included into the mechanism, the minimum of the voltage nullcline reflects opening of this current, and it is shifted away from the steep part of the Ca^2+^ nullcline (Fig. 4 B). This shifts the system away from the Andronov-Hopf bifurcation. Now, a downward shift of the voltage nullcline following inhibitory inputs moves its minimum across the flat part of the Ca^2+^ nullcline and produces a SNIC bifurcation. In this transition, the firing frequency gradually reduces to zero, and this allows a balance between NMDAR and GABAR conductances and restores the background firing frequency.

### Other studies of NMDA-GABA balance in DA neurons

The influence of NDMA and GABA receptor conductances on the DA neuron have been studied in several other papers [51–53]. The compensatory influence of NDMA and GABA receptor activation on the firing frequency has been predicted in modeling studies by Komendantov et al. [51]. Lobb et al. [53] modified our previous model [54] to capture the balance of the inhibition and excitation and disinhibition bursts. In these models, a high maximal frequency (>10 Hz) can be achieved by tonic activation of the AMPAR or in response to depolarizing current injection, which contradicts experimental observations [7,26]. To the contrary, a number of experimental studies suggest that stimulation of NMDA receptors evokes a burst of high-frequency firing, whereas AMPA receptor activation evokes modest increases in firing [29–32] (but see [43,55]). This is an important distinction, which impacts the excitability of the neuron. Further, in Komendantov et al. [51] and Canavier & Landry [52], the NMDAR conductances were restricted to dendrites, whereas GABAR conductance was somatic. The mechanism of frequency rise during dendritic application of NMDA is different from the mechanism of response to somatic NMDAR stimulation [54,56]. Somatic NMDAR stimulation has been shown to elicit high-frequency firing in earlier experiments [32,57] and used to achieve the NMDA-GABA balance [27]. Here, we base a new model on our previous model [56] that presented a mechanism for somatically-induced high-frequency firing in a reconstructed morphology first and reduced it to a single compartment. In the current model, we have integrated the mechanism for high-frequency firing together with the balance of NMDAR and GABAR activation.

### Intrinsic Ih current and synaptic inputs can switch the excitability type of the DA neuron

Changes in intrinsic currents can affect the excitability type and, thus, computational properties of the DA neuron. For instance, we observed that potentiation of Ih current promotes type II excitability of the simulated DA neuron. In addition to the contribution of Ih current to pacemaker activity, has been shown in DA neurons [38], as well as in the other neuronal types that Ih induces intrinsic subthreshold resonance [58,59,60]. Thus, augmentation of Ih current increases oscillatory behavior of the DA neurons, as well as their synchronization in response to excitatory pulses. However, low-frequency tonic firing could not be maintained at high conductances of Ih current, likely affecting background DA levels.

Further, the influence of tonic synaptic inputs can also change the transition to the rest state and, therefore, be described as altered excitability. Tonic activation of AMPA receptors or an elevated reversal potential of the GABAR conductance may make the low-frequency balanced state unreachable. The reason for that is a transition to type II excitability: firing is blocked at higher frequencies. Our explanation is that shunting is so strong that opening of the subthreshold sodium current cannot sustain the voltage growth. In other words, these changes unfold the voltage nullcline and bring its minimum close to the steep part of the Ca^2+^ nullcline (Fig. 5 B). This primes the system for the Andronov-Hopf bifurcation responsible for type II excitability. Further, we found that NMDAR activation also biases the neuron towards type II excitability. Although the type may change as parameters shift away from the boundaries of the firing region [61], together, these results suggest that in high-frequency regimes the DA neuron displays type II excitability. This switch in excitability type may play a role in a transient increase in DA concentration in response to salient stimuli as it is easier to synchronize type II neurons by an excitatory input. Figure 11 supports this hypothesis by showing higher transient DA release produced by heterogeneous population of DA neurons receiving synaptic AMPA and GABA tones than in the absence of synaptic tone. Thus, correlated excitatory synaptic inputs are more likely to evoke robust coincidence DA release when DA neurons display type II excitability. A growing body of literature links the type of excitability to neural coding [22,33,46,62,63,64,65]. For instance, Prescott et al. [66] suggested that type I neurons are best suited for coding stimulus intensity. Hence, the DA neuron is designed for encoding the intensity of the tonic depolarizing and hyperpolarizing inputs by its smooth frequency dependence. This further supports and augments a recently found unique computational role for the DA neuron: it performs subtraction of inhibitory and excitatory inputs [67]. The operation is optimal to calculate unpredicted value of an event, but rarely observed and hard to implement in the brain. The first type of DA neuron excitability is necessary to quantitatively encode the level of input by the firing rate and perform the subtraction. In conclusion, DA neurons can exhibit traits of both integrators and resonators and these traits are modulated by intrinsic and synaptic conductances. Depending on the current constitution, DA neurons can perform rate coding by integrating slow variations in the inputs and adjust basal DA concentration or they can detect transient coherent changes in the inputs and synchronize for producing robust DA transients.

## Methods

The biophysical model of the DA neuron is a conductance-based one-compartmental model:

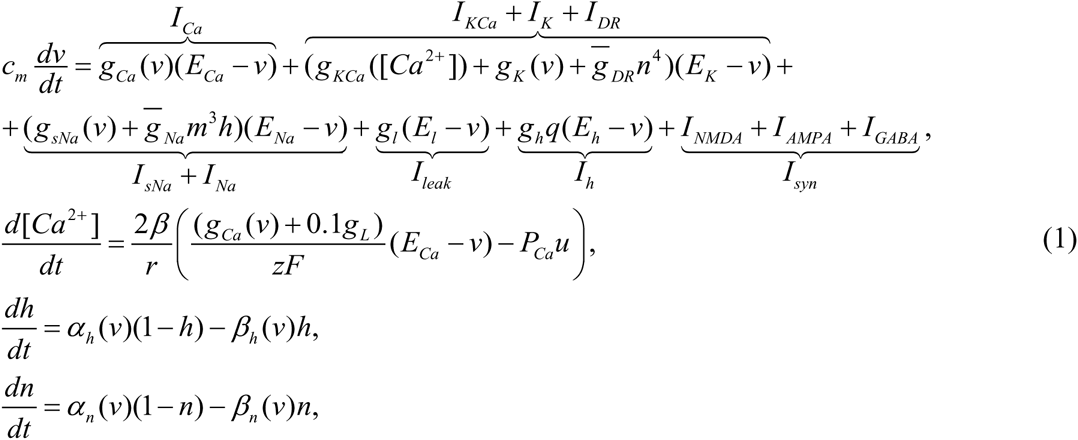
where *v* is the voltage, and *c*_*m*_ is membrane capacitance. There are eight intrinsic currents of the DA neuron: a calcium current (*I*_*Ca*_), a calcium-dependent potassium current (*I*_*KCa*_), a potassium current (*I*_*K*_), a direct rectifier current (*I*_*DR*_), a subthreshold sodium current (*I*_*sNa*_), a hyperpolarization-activated current (*I*_*h*_), a fast sodium current (*I*_*Na*_),and a leak current (*I*_*leak*_). The first subgroup of intrinsic currents: *I*_*Ca*_, *I*_*KCa*_, *I*_*K*_, *I*_*sNa*_, and *I*_*h*_ constitutes a pacemaking mechanism of the DA neuron. The second subgroup of the intrinsic currents (*I*_*Na*_, *I*_*DR*_) is responsible for spike generation. The last subgroup includes synaptic currents: the excitatory α-Amino-3-hydroxy-5-methyl-4-isoxazolepropionic (AMPA) and *N*-Methyl-D-aspartate (NMDA) receptor currents (*I*_*AMPA*_ and, *I*_*NMDA*_ respectively) and the inhibitory γ-Aminobutyric acid (GABA) receptor current (*I*_*GABA*_). Synaptic inputs can produce bursts and pauses in firing.

### Intrinsic oscillator

The main currents of the model that produce pacemaking activity of DA neuron are an L-type voltage-dependent calcium current (*I*_*Ca*_) and an SK-type calcium-dependent potassium current (*I*_*KCa*_). Gating of the calcium current is instantaneous and described by the function:

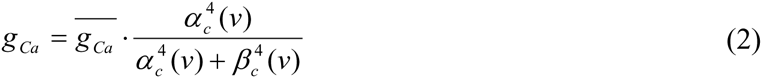

Calibration of the calcium gating function reflects an activation threshold of an L-type current, which is significantly lower in DA cells than in other neurons (˜ −50mV; [4]). Calcium enters the cell predominantly via the L-type calcium channel. Contribution due to the NMDA channel is minor [68]. Thus, calcium concentration varies according to the second equation of the system (1). It represents balance between Ca^2+^ entry via the L channel and a Ca^2+^ component of the leak current, and Ca^2+^ removal via a pump. In the calcium equation, *β* is the calcium buffering coefficient, i.e. the ratio of free to total calcium, r is the radius of the compartment, *z* is the valence of calcium, and *F* is Faraday’s constant. *P*_*Ca*_ represents the maximum rate of calcium removal through the pump. A large influx of Ca^2+^ leads to activation of the SK current, which contributes to repolarization as well as afterhypolarization of the DA cell. Dependence of the SK current (*I*_*KCa*_) on calcium concentration is modeled as follows

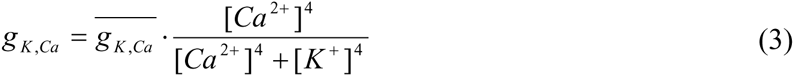

The neuron is repolarized by the activation of a large family of voltage-gated potassium channels. In addition to already described potassium currents, the model contains voltage-dependent K current (*I*_*K*_). Conductance of this current is given by a Boltzmann function:

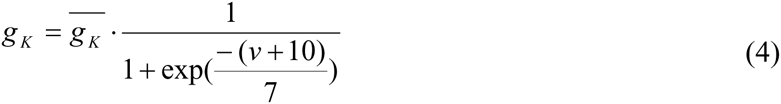

The DA neuron expresses voltage-gated sodium channels that carry a large transient current during action potentials (a spike-producing sodium current) and a noninactivating current present at subthreshold voltages (a subthreshold sodium current). Even though the persistent subthreshold sodium current is much smaller than the transient spike-producing current, it influences the firing pattern and the frequency of the DA neuron by contributing to depolarization below the spike threshold [50]. We modeled the voltage dependence of the subthreshold sodium current as follows

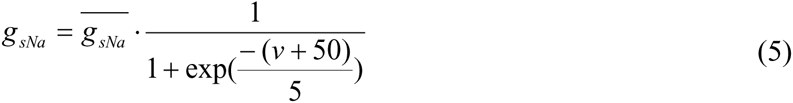

The kinetics and the voltage dependence of the subthreshold sodium current were taken from [69].

The majority of DA neuron express a hyperpolarization-activated nucleotide-gated (HCN) inward cation current (*I*_*h*_). The HCN current contributes to spontaneous firing of subpopulations of DA neurons [70]. The activation variable of *I*_*h*_ is governed by a first-order ordinary differential equation

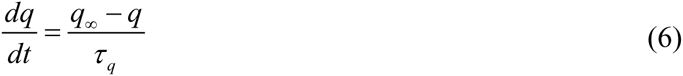

The maximal activation of *I*_*h*_ current is described by the following voltage-dependent equation [71]

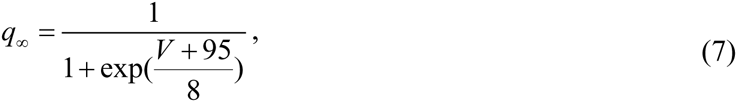

The voltage-dependent time constant is described by

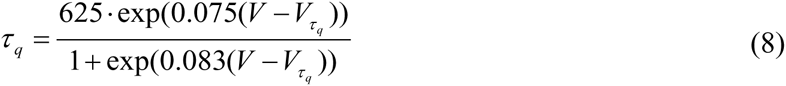

The leak current (*I*_*leak*_) in the model has the reversal potential of −35 mV, which is higher than in the majority of neuron types. In DA neurons, several types of depolarizing, nonselective cation currents are expressed, which likely contribute to depolarization during interspike intervals.

#### Model calibration

*Subthreshold currents*

Experiments show that subthreshold sodium currents contribute to depolarization towards the spike threshold [50,72]. Accordingly, the addition of the subthreshold sodium current into the model increases the background firing frequency of the DA neuron (data not shown). However, in the model this leads to a significant increase in maximal frequencies achieved during application of a constant depolarizing current or AMPA, whereas in experiments these frequencies have been shown to be limited by approximately 10 Hz [26]. We preserve this limit in the model by accounting for a Ca^2+^ component of the leak current. This reinforced the negative feedback loop through the Ca^2+^-dependent K^+^ current and limited the maximal frequencies. NMDAR activation increases the frequency, and our calibration of the model allowed an inhibitory or hyperpolarizing input to reduce the frequency to an arbitrary low value. Ca^2+^ entry through NMDA receptor was omitted to implement the spatial segregation of NMDAR and SK channels as in the previous models [51,56].

#### Spike-producing currents

We included spike-producing sodium current with maximal conductance 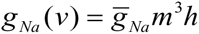 and potassium delayed rectifier current with maximal conductance 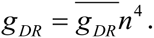. The activation of the Na^+^. current is assumed instantaneous and described by the function

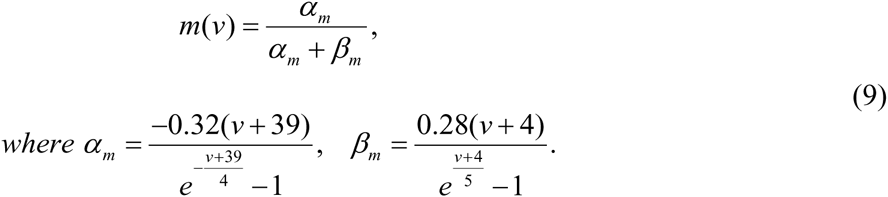

The inactivation kinetics is described by the equation for *h*in System (1), where

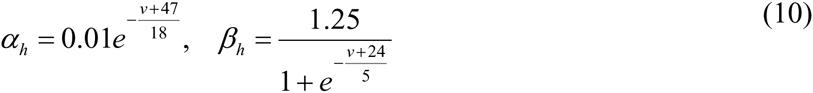

The delayed rectifier activation variable *n* is described by the 4th equation in System (1), where

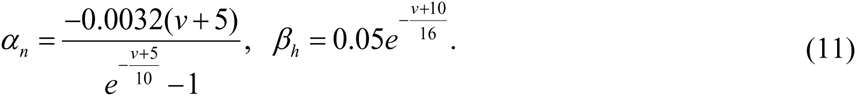

The currents are calibrated to produce a spike per each maximum of the voltage oscillations produced by the pacemaking mechanism without significantly changing the firing rate or pattern.

#### Synaptic inputs

DA neurons receive glutamatergic (Glu) excitatory drive through AMPA and NMDA receptors and inhibitory drive through GABA receptors. Changes in the membrane potential induced by synaptic conductances are described by the following equation

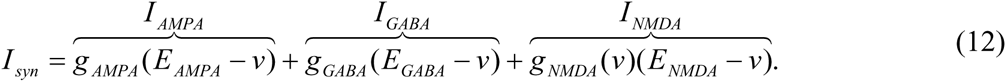

The AMPA and GABA conductances are voltage-independent, but the NMDA conductance has voltage sensitivity as in [19]

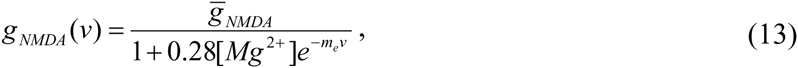

where [*Mg*^2+^] denotes the amount of magnesium, taken to be 0.5μM. The low slope of the voltage dependence (*m*_*e*_ =0.062) is critical for the increase in the frequency of spikes or subthreshold oscillations during NMDA application [56]. The receptors were activated tonically to mimic long-lasting injection of the conductances in dynamic clamp experiments [27,28,57], or iontophoresis of the agonists [31,32], or bath application of the agonists.

#### Modeling of Glu and GABA asynchronous inputs to the DA neuron

*Glu input*

Asynchronous Glu input to the DA neuron was produced by 35 Poisson distributed spike trains with frequencies of approximately 10 Hz. The number of spike trains was chosen to produce a relatively constant level of NMDA receptor activation, and, at the same time, take into account the effects of convergent synaptic inputs on the DA neuron, by thresholding NMDAR to activate only by coincidence of two or more spikes. The activation of the receptors in response to a synaptic input is described by the following equation

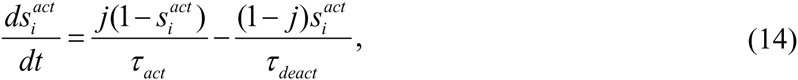

where *j* denotes a dimensionless synaptic input. It is normalized to change from 0 to 1 for 1 ms interval to mimic a single spike in the input. *i*denotes a receptor type, AMPA or NMDA. Desensitization of AMPA receptor is described by

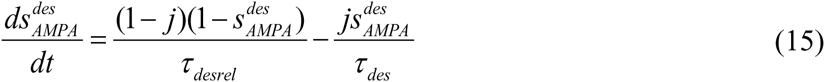

*GABA input:*

Asynchronous inhibitory input was produced by a population of 30 GABA neurons. The number of GABA neurons projecting to a DA neuron is not known. To choose this number, we note first that the percentage of GABAergic neurons varies between 12 and 45% in different subregions of the VTA [73]. Since the GABA neurons powerfully modulate DA neuron activity via direct, monosynaptic inhibitory connections [74,75,76,77,78], one can expect multiple GABA neurons to make connections with a single DA neuron. Second, we found that our results are valid for a wide range of the number of GABA neurons and start to change only if the number becomes small (10 or lower).

Voltage dynamics of each GABA neuron is described by the Wang-Buszaki equations of a fast spiking neuron [79],

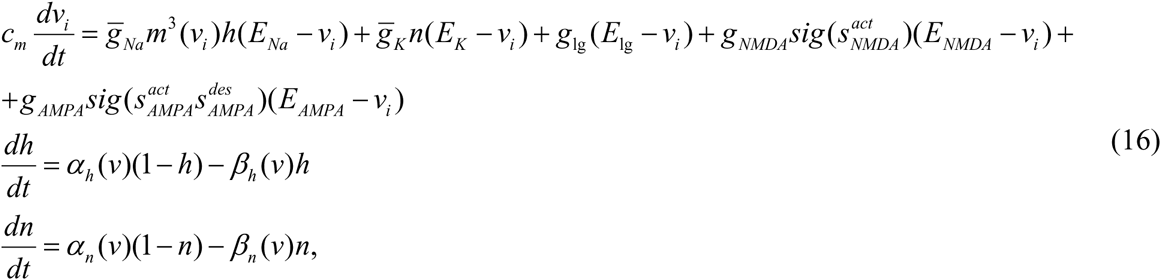

where *v*_*i*_ is a voltage of *i*_*th*_ GABA neuron in a population. The parameters of these equations were calibrated according to experimental observations [26]. Intrinsic firing frequencies of these neurons were set to a range of 12-22 Hz, similar to what is observed experimentally [26,80,81]. The activation/deactivation times of NMDA and AMPA receptors are the same for GABA as for DA neurons. The activity of each GABA neuron contributes to a GABAR current entering DA neuron, by contributing to activation of the gating variable according to the equation

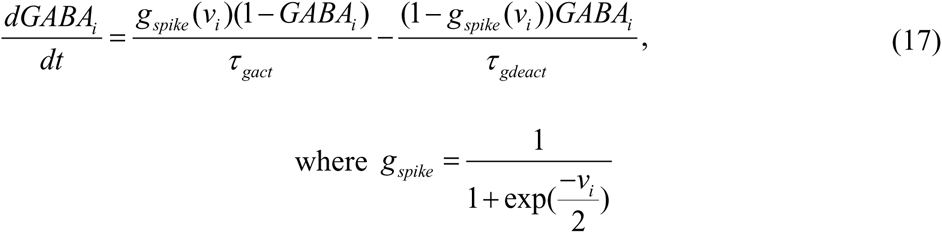

The total receptor activation is a sum of contributions produced by all GABA neurons in a population. A parameter that scales the GABA current is GABAR conductance *g*_*GABA*_. We normalize the GABA gating variable by the number of neurons in order to keep its value in a range between zero and one. Thus, for an asynchronous GABA population, the GABAR will be partially activated and the gating variable will have a low value. Model parameters are given in Table 1.

#### Model of DA release

The model of DA release is adopted from Wightman and Zimmerman (1990) [82] and is described by the following equation

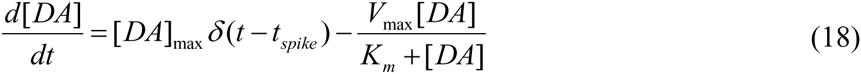

The first term describes the release from spiking activity of the DA neuron. Dirac delta function *δ* (*t-t*_*spike*_) represents the release at time of a spike. Maximum amount of DA released per spike is [*DA*]_max_ = 0.1 *μM*. The second term represents DA uptake described by Michaelis-Menten equation, where *V*_*max*_ = 0.004μM/*ms* is the maximal rate of uptake by a transporter and *K*_*m*_ = 0.2μ*M* is the affinity of the transporter for dopamine.

#### Modeling a heterogeneous population of DA neurons and their inputs

Heterogeneity in the population of DA neurons was putatively introduced by varying the leak conductance. Further, neurons received correlated fluctuating NMDA inputs. NMDAR conductance to each DA neuron was given by a linear summation of Ortein-Uhlenbeck (OU) processes [83] described as following:

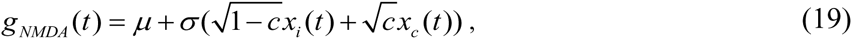

where *μ* = 1.5*mS* / *cm*^2^ and σ = 0.5*mS* / cm^2^ are the values of the mean and the standard deviation of the NMDAR conductance used for the simulations. *x*_*c*_(*t*) is the common component of the NMDA input that was applied to all of the DA neurons, whereas *x*_*i*_(*t*) is the independent component, which was generated individually for each neuron. A shared fraction of the input is determined by the input correlation *c*and was set to *c* = 0.5. Each OU process was formed by the following equation:

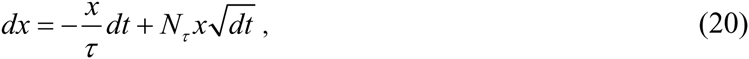

where *x*(*t*) is Gaussian white noise with zero mean and unit variance. *N*_*τ*_ = (2 / τ)^½^ is a normalization constant that makes *x*(*t*) have unit variance. A correlation time *τ* = 5*ms* was used [24,84].

#### Spike detection and firing pattern quantification in the model

In the model with fast sodium and the delayed rectifier potassium spike-producing currents, a spike was registered whenever voltage oscillation reached the threshold of 0 mV. In the reduced model (without spike-producing currents), a spike was registered every time voltage oscillations crossed the threshold of −40 mV, as experimentally it was shown that a DA neuron action potential is triggered when the voltage is depolarized to approximately −40 mV [3]. Voltage oscillations that were below these thresholds in the models with and without the spike-producing currents respectively were not counted as spikes and did not contribute to the firing frequency. To analyze firing pattern of simulated DA neuron in the presence of different synaptic currents, we quantified its firing rate and bursting. Mean firing rate of the simulated DA neuron was calculated as an inverse of the mean interspike interval (ISI). To calculate bursting we used ISI coefficient of variation (CV), calculated as the SD/mean of 200 ISIs.

## Acknowledgments

Authors thank Dr. Oleg Morozov, Maxym Myroshnychenko and Matteo di Volo for their support and constructive discussions.

**Table 1.** Model parameters

| Parameter | Description | Value |
| --- | --- | --- |
| $c_m$ | Membrane capacitance | $1\mu F / cm^2$ |
| $\bar{g}_K$ | Potassium conductance | $1mS / cm^2$ |
| $g_{Na}$ | Sodium conductance | $50 mS/cm^2$ |
| $g_{DR}$ | Conductance of delayed rectifier | $2 mS/cm^2$ |
| $\bar{g}_{Ca}$ | Calcium conductance | $2.5mS / cm^2$ |
| $\bar{g}_{KCa}$ | Calcium-dependent potassium conductance | $7.8mS / cm^2$ |
| $\bar{g}_{sNa}$ | Subthreshold sodium conductance | $0.13mS / cm^2$ |
| $g_{leak}$ | Leak conductance | $0.18mS / cm^2$ |
| $E_K$ | Potassium reversal potential | $-90mV$ |
| $E_{Ca}$ | Calcium reversal potential | 50mV |
| $E_{Na}$ | Sodium reversal potential | 55mV |
| $E_h$ | HCN reversal potential | -20mV |
| $E_{leak}$ | Leak reversal potential | -35mV |
| $E_{NMDA}$ | NMDA reversal potential | 0mV |
| $E_{AMPA}$ | AMPA reversal potential | 0mV |
| $E_{GABA}$ | GABA reversal potential | -90mV |
| $\tau_{aact}$ | AMPA receptor activation time | 1ms |
| $\tau_{adeact}$ | AMPA receptor deactivation time | 1.6ms |
| $\tau_{ades}$ | AMPA receptor desensitization time | 6.1ms |
| $\tau_{adesrel}$ | AMPA receptor release from desensitization time | 40ms |
| $\tau_{nact}$ | NMDA receptor activation time | 7ms |
| $\tau_{ndeact}$ | NMDA receptor deactivation time | 170ms |
| $\tau_{gact}$ | GABA receptor activation time | 0.08ms |
| $\tau_{gdeact}$ | GABA receptor deactivation time | 10ms |
|  |  | 10ms |

